## Supporting Information for "Mechanism of Dimer Selectivity and Binding Cooperativity of BRAF Inhibitors"

[1]Department of Pharmaceutical Sciences, University of Maryland School of Pharmacy, Baltimore, MD 21201, United States; [2]Division of Applied Regulatory Science, Office of Clinical Pharmacology, Office of Translational Sciences, Center for Drug Evaluation and Research, United States Food and Drug Administration, Silver Spring, MD 20993, United States; [3]Department of Biochemistry, Department of Medicine, Department of Oncology, Montefiore Einstein Comprehensive Cancer Center, Albert Einstein College of Medicine, New York, NY 10461, United States; [4]Department of Oncological Sciences, Icahn School of Medicine at Mount Sinai, New York, NY 10029, United States

### Supplemental tables

| Inhibitor | PDB | $\alpha$C;DFG | BPs | BP H-bonds | $\alpha$C position | K-E |
|---|---|---|---|---|---|---|
| | | | **Monomer selective** | | | |
| Vemurafenib | 5JRQ | CODI | I,II | D594, G596 | 21.5/21.5 | 11.5/11.8 |
| Dabrafenib | 5CSW | CODI | I,II | K483, D594, G596 | 21.4/21.8 | 7.4/8.7 |
| PLX7904 | 4XV2 | CODI | I,II | D594, F595, G596 | 20.9/20.4 | 7.5/8.7 |
| | | | **Equipotent** | | | |
| AZ628 | 4G9R | CIDO | I,II,III | E501, D594 | 19.5/19.5 | 3.0/2.6 |
| TAK632 | 4KSP* | CIDO | I,II,III | E501, D594 | 18.8/19.0 | 3.0/3.9 |
| LY3009120 | 5C9C | CIDO | I,II,III | E501, D594 | 19.2/19.2 | 2.7/2.7 |
| Ponatinib | 6P3D | CIDO | I,II,III | E501, D594<br>I573, H574 | 19.2/19.2 | 3.0/3.0 |
| Lifirafenib (BGB283) | 4R5Y | CIDO | I,II,III | E501 | 19.0/18.7 | 5.5/5.1 |
| Tovorafenib (TAK580) | 6V34 | CIDO | I,II,III | E501, D594 | 18.6/18.8 | 2.5/2.9 |
| SB5909885 | 2FB8* | CIDI | I | E501 | 18.6/18.3 | 2.9/3.2 |
| GDC0879 | 4MNF | CIDI | I | E501 | 18.4/18.7 | 2.8/2.9 |
| | | | **Dimer selective** | | | |
| Naporafenib (LXH254) | 8F7P* | CIDO | I,II,III | E501, D594 | 19.3/19.3 | 3.0/3.0 |
| RAF709 | 5VAM* | CIDO | I,II,III | E501, D594 | 19.2/19.1 | 2.8/2.8 |
| Sorafenib | 1UWJ | CIDO | I,II,III | E501, D594 | 18.8/18.8 | 2.6/2.6 |
| Belvarafenib | 6XFP* | CIDO | I,II,III | E501, D594 | 19.3/19.3 | 2.7/2.7 |
| PHI1 | 6P7G | CIDO | I,II,III | E501, D594, H574 | 19.1/19.0 | 2.5/2.5 |

**Table 1.** List of the $BRAF^{V600E}$ inhibitors and the structure features of the co-crystal structures in the PDB. The monomer and dimer selectivities of inhibitors in black are based on the experimental data in Ref (***Adamopoulos et al., 2021***) and Ref (***Cotto-Rios et al., 2020***) (PHI1). The monomer and dimer selectivities of inhibitors in red were predicted by us and supported by experiments in Ref (***Karoulia et al., 2016***) (GDC0879) and Ref (***Tkacik et al., 2023***) (Tovorafenib). Note, the PDB entries indicated by an astrisk are co-crystal structures in complex with the wild type BRAF (the $BRAF^{V600E}$ forms are unavailable). All structures contain an inhibitor in each protomer, with the exception of PLX7904; in the PDB entry 4XV1, PLX7904 is only present in protomer A. The $\alpha$C helix position (in Å) is defined in the main text. Two values refer to the two protomers. The back pockets (BPs) occupied was calculated by KLIFS (***Kooistra et al., 2016***) based on the definition of Liao (***Liao, 2007***). K-E refers to the distance (in Å) between the amine nitrogen of Lys483 and the nearest carboxylate oxygen of Glu501.

| System, protomer | $\alpha$C position (Å) | K-E distance (Å) | DFG dihedral (°) |
|---|---|---|---|
| Apo monomer, A | 23.3 ± 1.7 | 8.1 ± 3.1 | 214 ± 19 |
| Apo dimer, A | 22.2 ± 1.0 | 6.5 ± 2.9 | 250 ± 39 |
| Apo dimer, B | 22.0 ± 1.7 | 5.5 ± 2.2 | 201 ± 24 |
| PHI1 (mixed), apo | 21.3 ± 1.0 | 5.5 ± 3.1 | 239 ± 41 |
| PHI1 (mixed), holo | 18.9 ± 0.7 | 2.8 ± 0.3 | 283 ± 24 |
| LY (mixed), apo | 22.8 ± 1.7 | 5.3 ± 2.4 | 204 ± 35 |
| LY (mixed), holo | 19.0 ± 0.5 | 4.9 ± 1.2 | 295 ± 13 |
| PHI (holo), A | 18.3 ± 0.6 | 2.8 ± 0.1 | 285 ± 17 |
| PHI (holo), B | 18.7 ± 0.7 | 2.8 ± 0.1 | 303 ± 12 |
| LY (holo), A | 19.4 ± 0.6 | 3.4 ± 1.1 | 301 ± 18 |
| LY (holo), B | 19.2 ± 0.5 | 4.3 ± 1.4 | 295 ± 13 |

**Table 2.** Average value and standard deviation of reported quantities from out dimeric BRAF simulations, separated by system and protomer. Each quantity was calculated for each replica after removing the first 2 $\mu$s for equilibration.

### Supplemental Figures

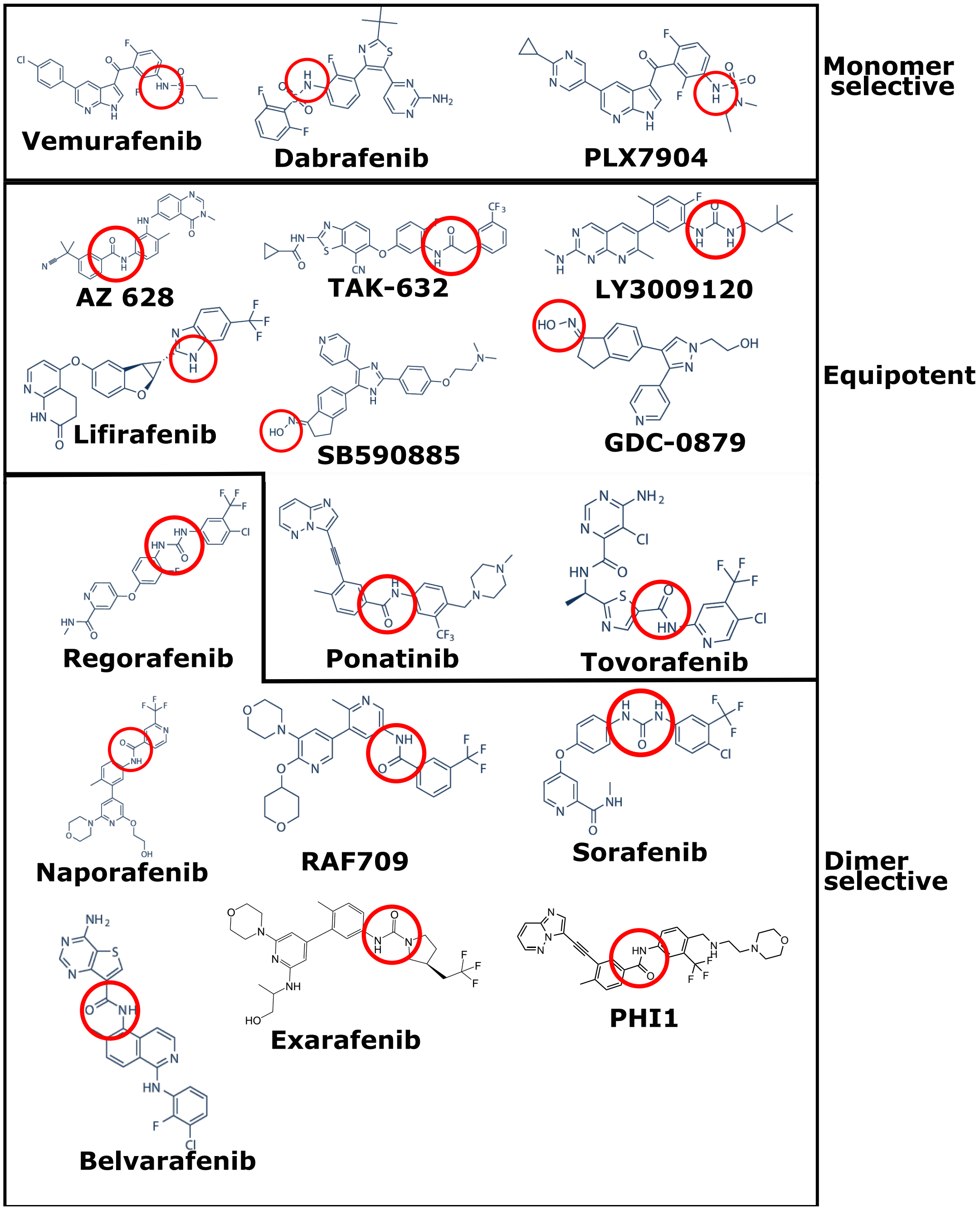


**Figure 1.** Chemical structures of the monomer-selective, equipotent, and dimer-selective BRAF$^{V600E}$ inhibitors in Supplementary Table 1. The chemical group responsible for forming the hydrogen bonds with E501 and/or D594 are circled in red.

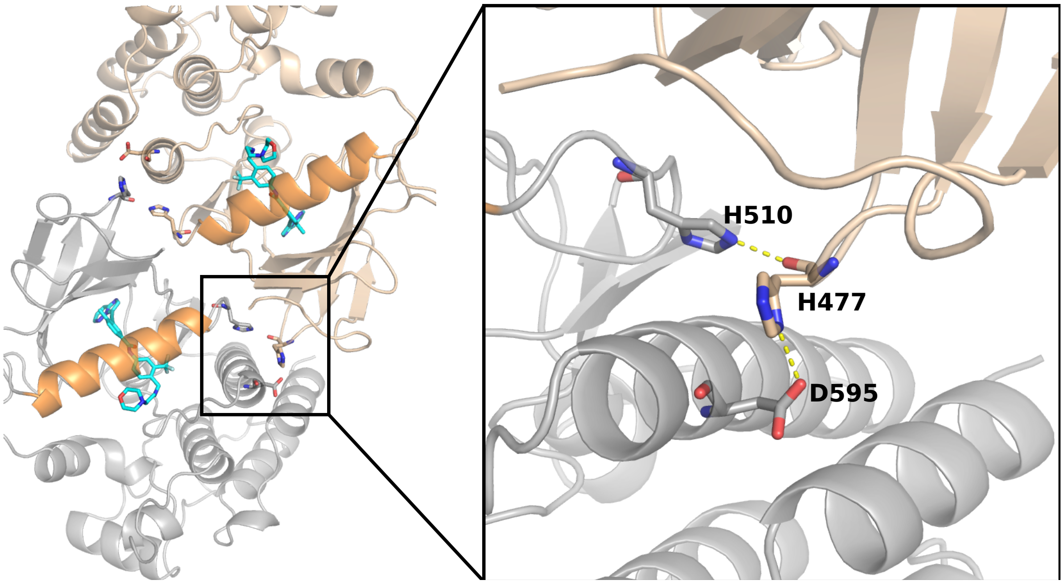


**Figure 2. Visualization of the H510-H477-D595 interactions at the dimer interface of BRAF$^{V600E}$. Left.** Visualization of the BRAF$^{V600E}$ dimer (PDB ID: 6P7G) (***Cotto-Rios et al., 2020***). The $\alpha$C helix is colored brown and PHI1 is colored cyan. The dimer interface residues H477, H510, and D595 are shown in sticks. **Right.** A zoomed-in view of the interactions between H477 in protomer A (light brown color) and H510 or D595 in the protomer B (grey color).

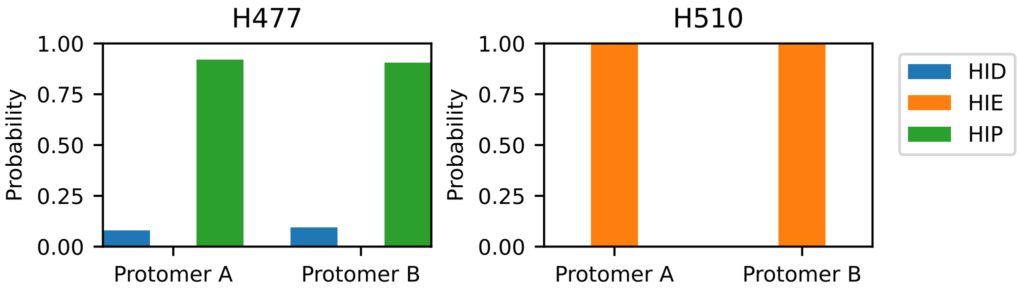


**Figure 3. Protonation and tautomer states of the interface histidines determined by the all-atom PME-CpHMD titration. Left.** Calculated probabilities of the neutral (HID in blue, HIE in orange) and charged (HIP in green) states for H477 (**left**) and H510 (**right**) at pH 7.5. The probabilities of protonation states for either histidine is nearly identical between the two protomers. As a default in the CpHMD program ***Harris et al.*** (***2022***), the charged HIP state was defined by $\lambda < 0.2$, while the neutral states were defined by $\lambda > 0.8$ and $\chi < 0.2$ (HID) or $\chi < 0.8$ (HIE). The last 5 ns of the replica runs were used for the calculation.

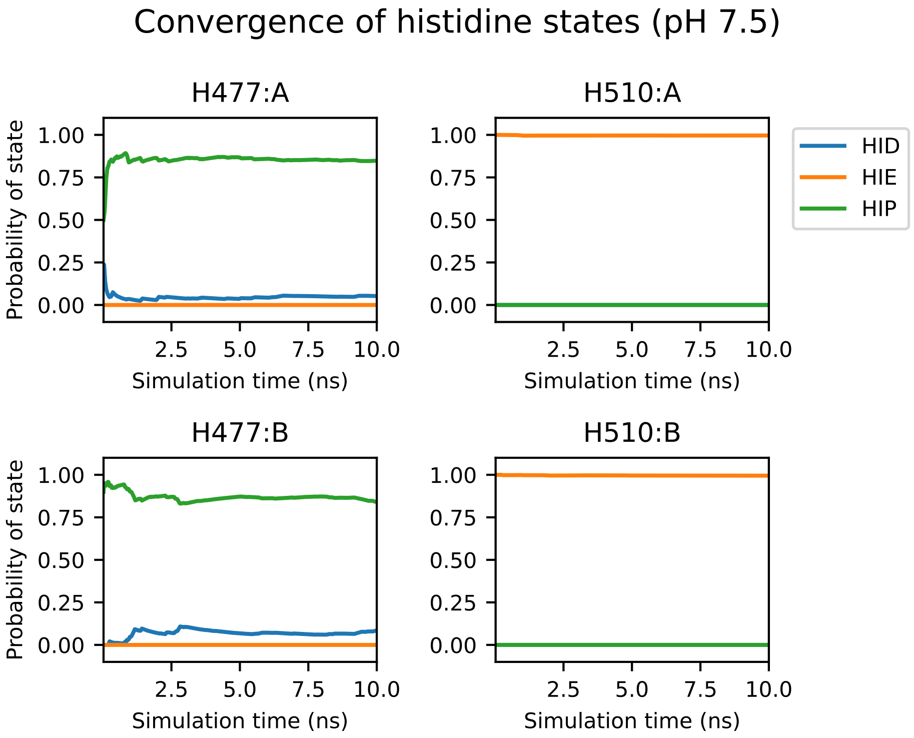


**Figure 4. Convergence of the protonation/tautomer states of H477 and H510 at pH 7.5.** Running average probability of each protonation/tautomer state for H477 and H510 at pH 7.5 as a function of simulation time (top plots for protomer A and bottom plots for protomer B). Convergence of the probability occurs after roughly 2 ns, with little change after 5 ns.

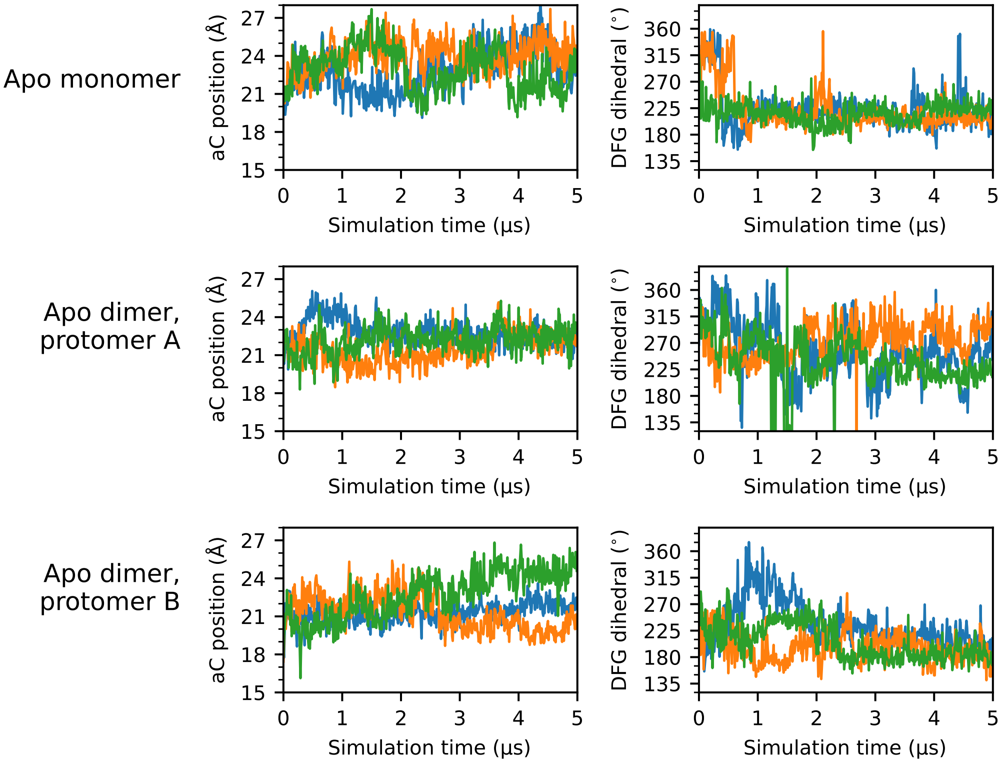


**Figure 5. Time series of $\alpha$C-helix position and DFG dihedral for all apo simulations.** Time series plots of the $\alpha$C-helix position (left column) and DFG dihedral (right column) for each apo protomer simulated. Each of the three replicas are represented as a separate line; for simplicity, a stride of 10 ns was used while plotting.

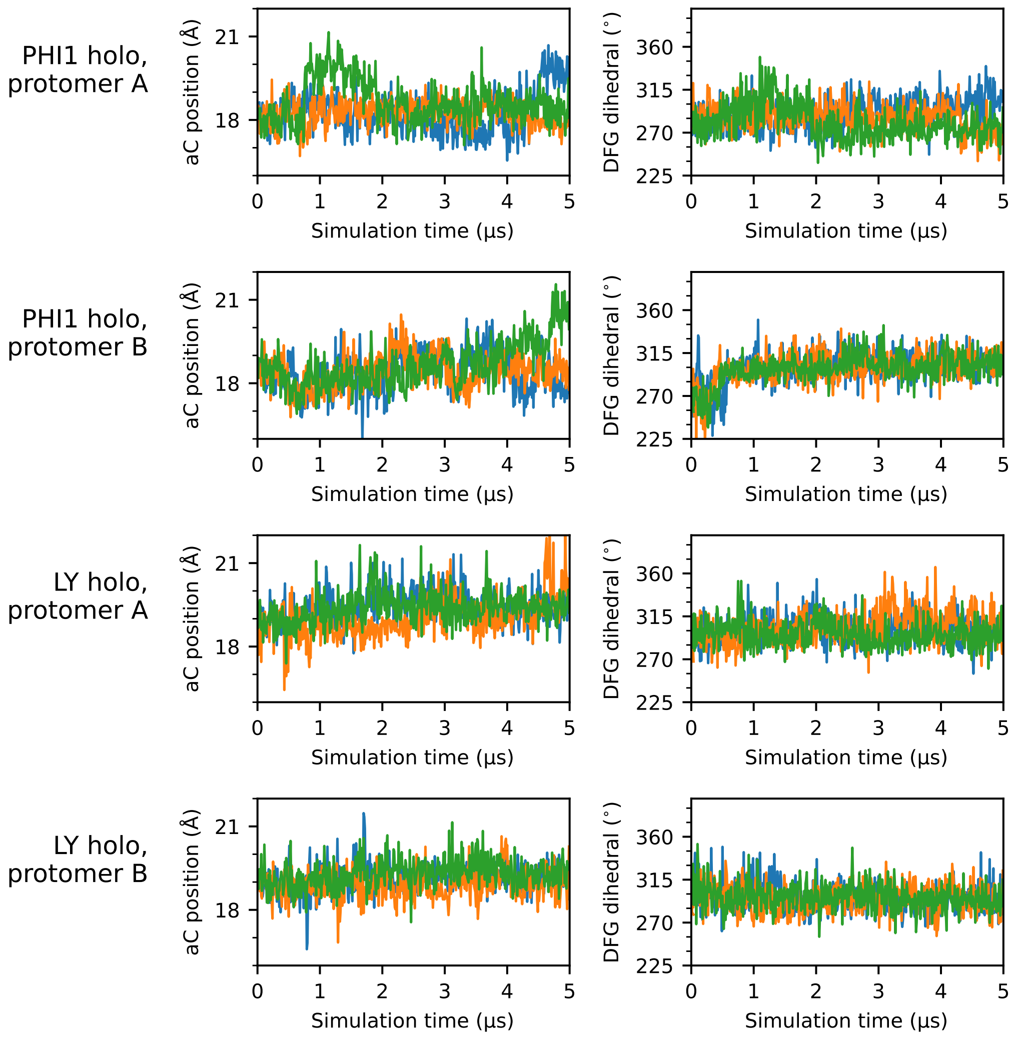


**Figure 6. Time series of $\alpha$C-helix position and DFG dihedral for all holo simulations.** Time series plots of the $\alpha$C-helix position (left column) and DFG dihedral (right column) for each holo protomer simulated. Each of the three replicas are represented as a separate line; for simplicity, a stride of 10 ns was used while plotting.

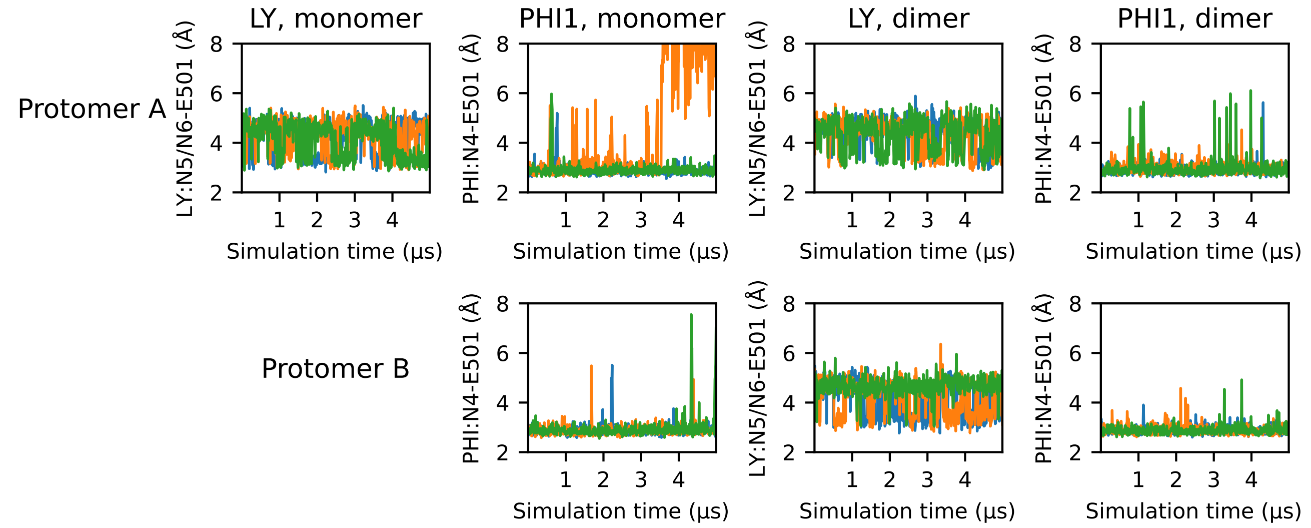


**Figure 7. Time series of E501-ligand hydrogen bonding.** Time series of the minimum distance between E501 carboxylate oxygens and the proton donor of LY (N5 or N6) and PHI1 (N4). Each of the three replicas are represented as a separate line; for simplicity, a stride of 10 ns was used while plotting.

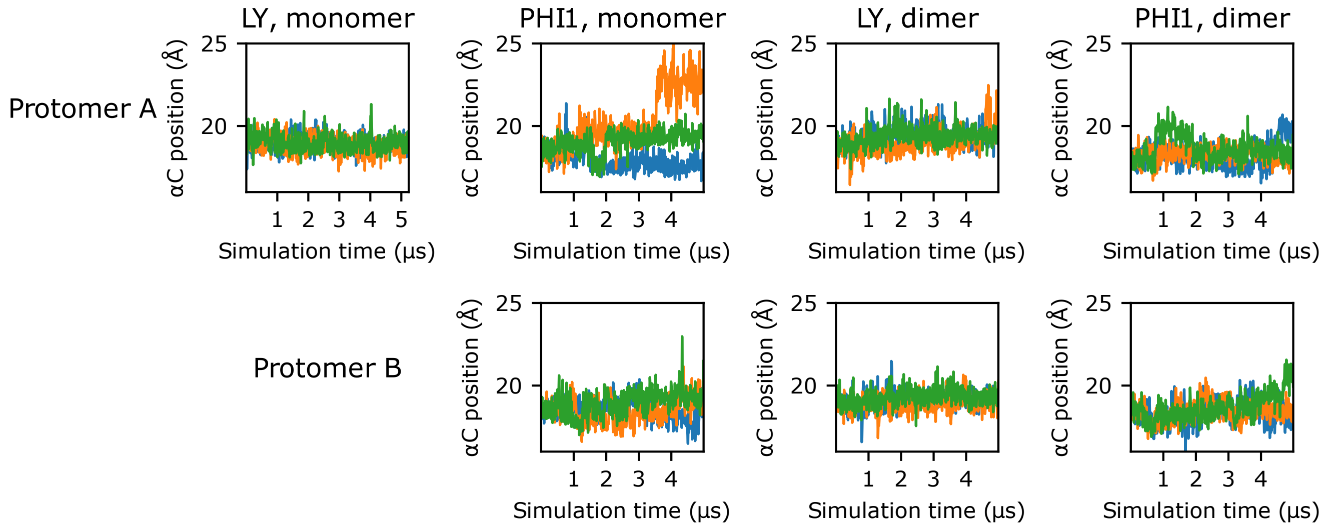


**Figure 8. Time series of $\alpha$C-helix position.** Time series of the $\alpha$C-helix position for inhibited monomer and dimer simulations. Each of the three replicas are represented as a separate line; for simplicity, a stride of 10 ns was used while plotting.

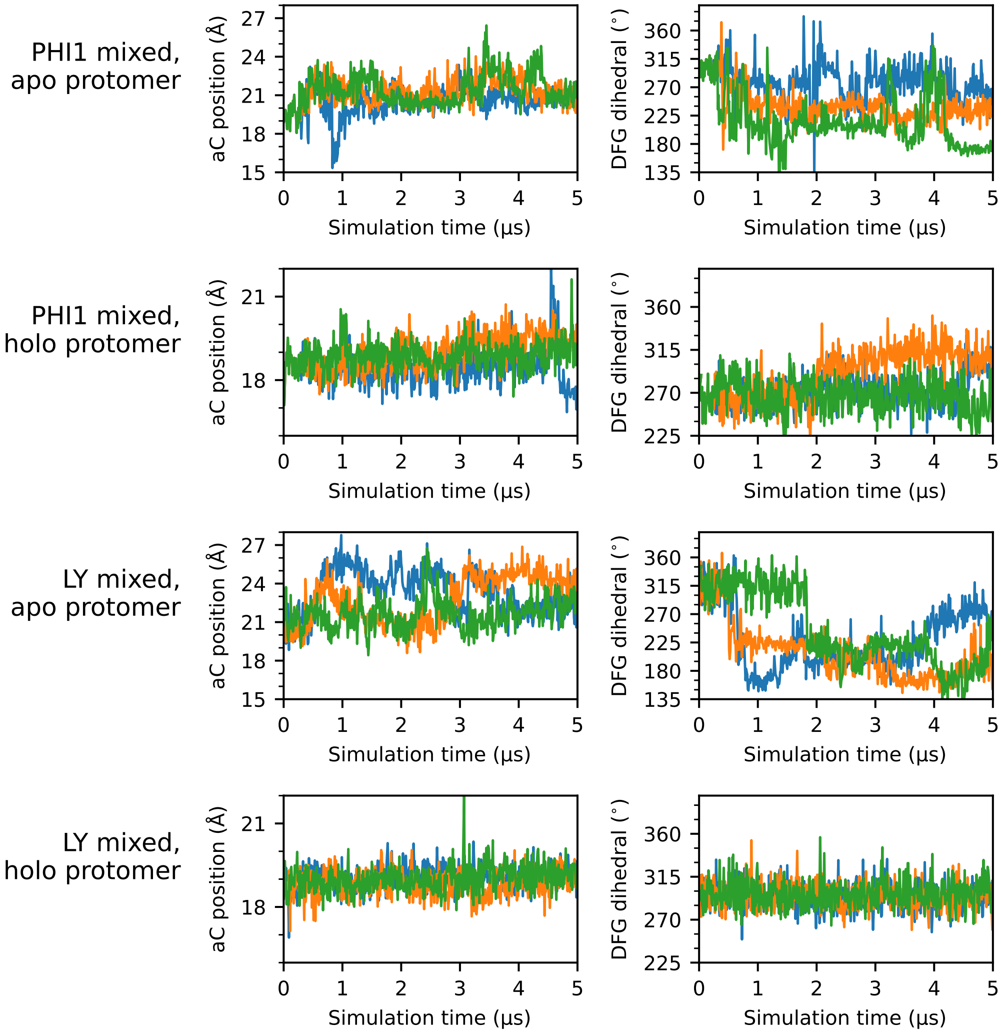


**Figure 9. Time series of $\alpha$C-helix position and DFG dihedral for all mixed simulations.** Time series plots of the $\alpha$C-helix position (left column) and DFG dihedral (right column) for each protomer simulated in the mixed simulations. Each of the three replicas are represented as a separate line; for simplicity, a stride of 10 ns was used while plotting.

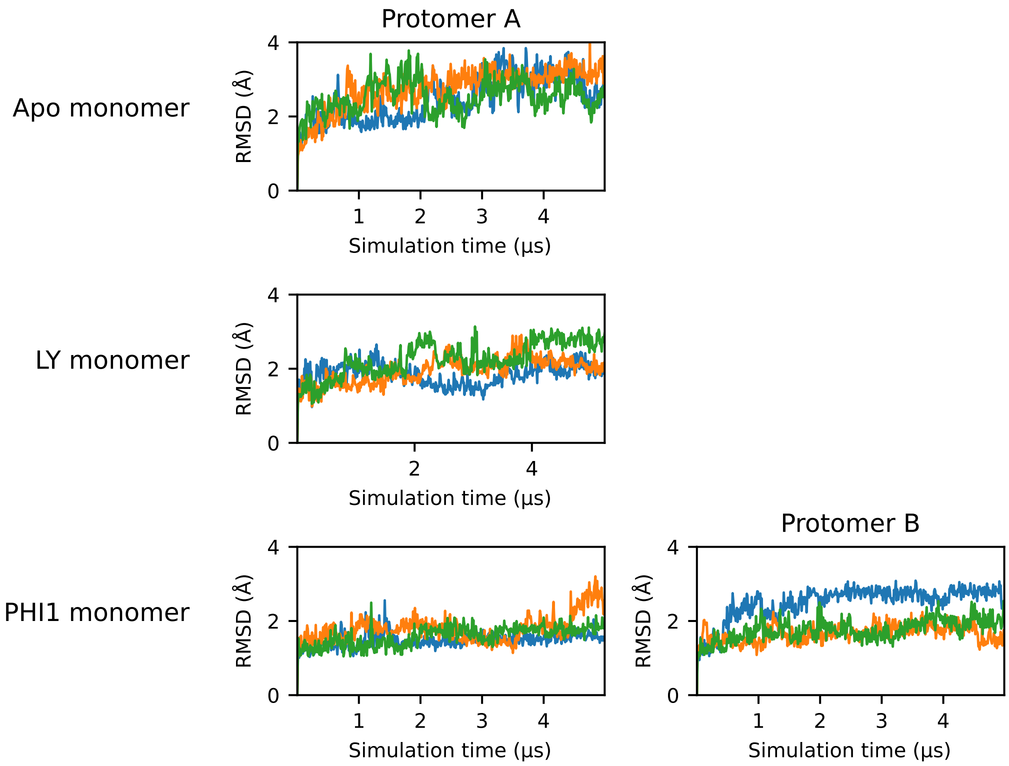


**Figure 10. RMSD time series for the monomer simulations.** Time series plots of the root-mean-square deviation (RMSD) of the protomer in each monomer simulation. The RMSD was calculated using heavy atoms of the protomer backbone, excluding residues on the a-loop and $\alpha$C-helix. Each of the three replicas are represented as a separate line; for simplicity, a stride of 10 ns was used while plotting.

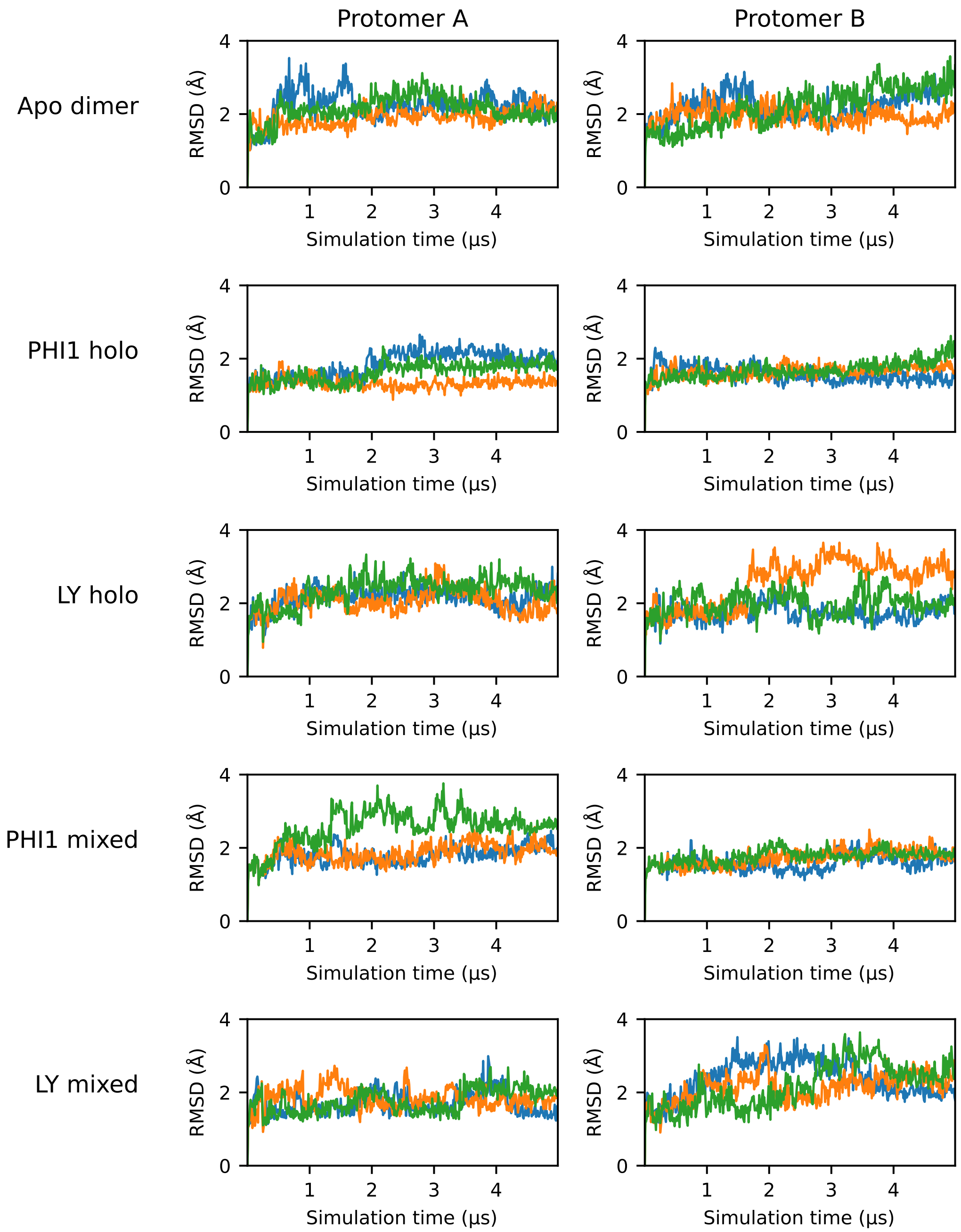


**Figure 11. RMSD time series for the dimer simulations.** Time series plots of the RMSD of the protomer in each dimer simulation. The RMSD was calculated using heavy atoms of the protomer backbone, excluding residues on the a-loop and $\alpha$C-helix. Each of the three replicas are represented as a separate line; for simplicity, a stride of 10 ns was used while plotting.
